# Probiotic biogeography and sepsis prevention in the neonatal intestine

**DOI:** 10.1101/2025.09.26.678848

**Authors:** Sierra C. Hansen, Christopher W. Hamm, Casey T. Weaver, Michael J. Gray

## Abstract

Neonatal infection is one of the leading causes of neonatal morbidity and mortality worldwide, particularly in those born prematurely or with low birth weight. Probiotic bacteria have been demonstrated to protect against the development of neonatal intestinal dysbiosis and are widely used in peri- and post-natal clinical settings. However, formulations and efficacy are highly variable, highlighting a critical gap in the current understanding of the mechanistic underpinnings of successful probiotic interventions in this population. Furthermore, current studies on probiotic efficacy largely rely on indirect or relative readouts of intestinal bacterial burden. Herein, we directly mapped the biogeography of intestinal colonization and quantify the probiotic effects of *Escherichia coli* Nissle 1917 (EcN) and *Ligilactobacillus murinus* strain V10 against *Klebsiella pneumoniae* dysbiosis across the span of the neonatal murine intestine. Despite substantial differences in biogeography within the intestine, both EcN and *L. murinus* V10 significantly reduced *K. pneumoniae* colonization and mortality from *K. pneumoniae* sepsis, with EcN doing so much more robustly. EcN’s probiotic effect was partially dependent on its ability to respire oxygen. Contrary to the dominant paradigm and practice in the probiotic field, combining multiple probiotic strains did not necessarily increase efficacy. Simultaneous treatment with EcN and *L. murinus* V10 was less effective than EcN treatment alone at preventing death from sepsis. These results highlight important variables which must be taken into account in the design of effective future probiotic intervention strategies.

**IMPORTANCE:** In this work we use a mouse model of late-onset neonatal sepsis (LOS) to rigorously test fundamental assumptions that underlie the current paradigm for understanding the impact of probiotics on intestinal disease. We demonstrate that two distantly related probiotic bacteria (*Escherichia coli* Nissle 1917 and *Ligilactobacillus murinus* V10) can each effectively reduce both intestinal colonization and death caused by the LOS pathobiont *Klebsiella pneumoniae*, acting by distinct ecological and molecular mechanisms. Our results provide new evidence that will be critical for designing and implementing safe and effective probiotic treatment regimens for LOS, a devastating and difficult to treat disease. More broadly, our results show that ecological principles are key to understanding how interventions that modulate the gut microbiome work, and that some of the assumptions underlying current interventions need to be reevaluated, especially when it comes to combining multiple probiotic strains and species.

## INTRODUCTION

Neonatal infection remains a leading cause of both neonatal morbidity and mortality worldwide, particularly in preterm infants. Preterm infants are uniquely susceptible to the development of neonatal dysbiosis. One study found that premature membrane rupture and intra-amniotic infection impacts between 25-30% of preterm infants (1), and these events are known to result in dysbiosis (2). In contrast to term infants, delivery *via* cesarian section is both more common and typically considered the safer mode of delivery in this patient population (3, 4). Following delivery, many preterm infants require stabilization and often face prolonged stays in neonatal intensive care units (NICUs). This setting profoundly impacts both the acquisition of pioneering microbiome members and the initial trajectory of the gut microbiome and significantly increases the risk of nosocomial infection (5).

The most commonly cited short-term consequences of dysbiosis at this developmental stage are necrotizing enterocolitis (NEC), a necrotic infection of the large intestine (6), and sepsis, including late-onset sepsis (LOS), defined as sepsis presenting between 72 hours and 28 days after birth (7), Both NEC and LOS remain unfortunately common and difficult to treat causes of both morbidity and mortality in preterm infants, particularly among those with a low or very-low birthweight (8–10), While establishment of a causal relationship remains elusive, intestinal dysbiosis followed by translocation of intestinal bacteria into the bloodstream is the leading clinical hypothesis behind these disease states as studies have demonstrated that bacteria isolated from blood culture in septic infants matched those found in the gut microbiome (11–14), and increased presence and growth of beneficial bacteria has been demonstrated to protect against the development of sepsis in both animal (15) and human (13) studies.

Probiotic bacteria have been demonstrated to protect against the development of neonatal intestinal dysbiosis (16–23). However, probiotic formulations and efficacy vary substantially, highlighting a critical gap in the current understanding of the mechanisms governing probiotic efficacy in this population. This gap is exacerbated by regulatory standards which favor economic expansion over scientific rigor. In addition, in 2023, the US Food and Drug Administration issued a blanket warning against the use of probiotics in preterm infants following a single case report of a preterm infant that developed sepsis following administration of the probiotic strain *Bifidobacterium longum* subsp. *infantis* EVC001 and subsequently died (24). This warning has greatly reduced the use of probiotics in US neonatal care units. Important to note, however, is that details on the cause of death and concomitant diseases or conditions for this individual were not provided. The sweeping implications and impact of this warning have been publicly criticized by multiple groups and organizations (25, 26) including the European Society for Paediatric Gastroenterology, Hepatology and Nutrition (ESPGHAN) and the European Foundation for the Care of Newborn Infants (EFCNI)(26), who argue that probiotics have an important and effective place in the management of neonatal health.

In this work, we directly quantified the probiotic effects of two very different probiotic bacteria, *Escherichia coli* Nissle 1917 (EcN 1917) and *Ligilactobacillus murinus* V10, against the pathobiont *Klebsiella pneumoniae* in a murine model of late-onset neonatal sepsis. The results of these experiments demonstrated that specific probiotic interventions can effectively reduce both dysbiosis and death from sepsis, that different probiotics act in different intestinal compartments and by different ecological and likely by different molecular mechanisms, and that, contrary to the dominant paradigm in the field, combining multiple probiotics does not necessarily lead to increased protective effects.

## RESULTS

Much of the intestinal development that takes place *in utero* in humans occurs postnatally in rodents (27), meaning that the newborn mouse is more similar developmentally to a preterm than a full-term human infant. Similarly to preterm human infants, neonatal mice exhibit delayed appearance of obligate anaerobes (15). To model dysbiosis and late-onset sepsis, we therefore gavage neonatal (5-7-day old, depending on the experiment) mice with specific strains of *Klebsiella pneumoniae* (15). We used direct plate counting to determine the actual viable bacterial loads in the intestines of neonatal mice treated with *Kp-39*. It is important to note that strain *Kp-39* does not cause sepsis in neonatal mice (15), so experiments with this strain specifically test the ability of probiotic treatments to reduce dysbiotic overgrowth of *K. pneumoniae* in the intestine. This is in contrast to the more virulent *K. pneumoniae* strain *Kp-43816*, which leads to fulminant sepsis and death in our model (see below)(15).

### EcN 1917 reduces *Kp-39* colonization burden across the intestinal span

Probiotic administration of EcN 1917 (Fig. 1A) significantly reduced *Kp-39* colonization on a total intestinal level (Fig. 1B), which can be attributed to significant reductions in each of the four intestinal segments examined (Fig. 1C).

**Figure 1.**
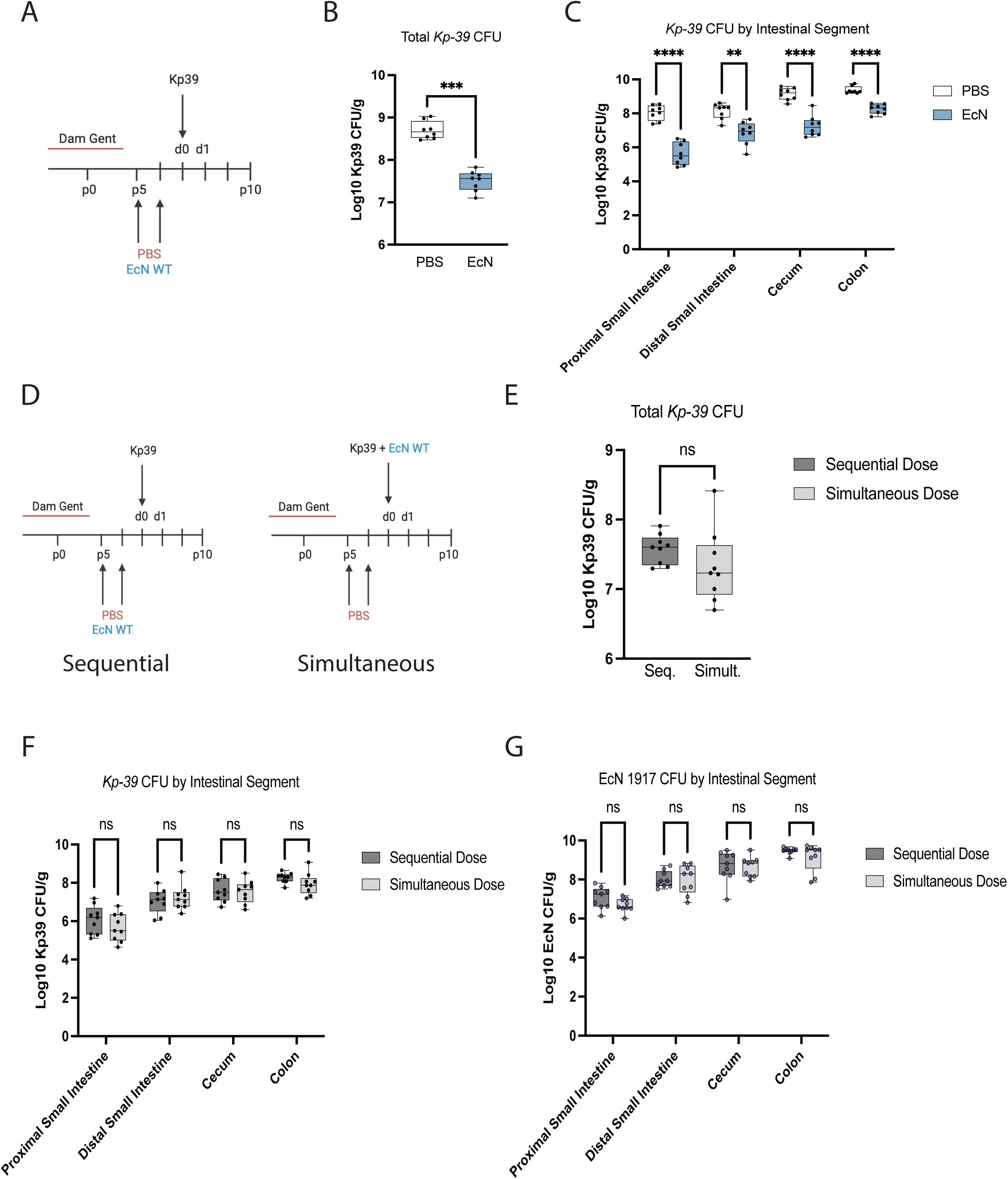
Biogeography of EcN 1917 probiotic effect in the neonatal intestine. A, Schematic illustration of murine model of neonatal sepsis with probiotic treatment. Littermate pups from gentamicin-treated dams were treated i.g. on day 5 (P5) and 6 (P6) of life with either EcN 1917 or PBS control before infection with 10^7^ CFU *Kp-39* on day 7 (P7). B, Total intestinal *Kp-39* colonization burden by treatment group on treatment day 1 (d1). C, *Kp-39* colonization by treatment group and intestinal segment on d1. Data were pooled from two independent experiments of within-litter controls with PBS (*n* = 8) and EcN (*n* = 8). D, Schematic illustration of neonatal priority effect model. Littermate pups from gentamicin-treated dams were treated i.g. on day 5 (p5) and 6 (p6) of life with either EcN 1917 (sequential dose) or PBS control (simultaneous dose) before infection with 10^7^ CFU *Kp-39* (sequential dose) or 10^7^ CFU *Kp-39* and EcN 1917 (simultaneous dose) on day 7 (P7). E, Total intestinal *Kp-39* colonization burden by treatment group on treatment day 1 (d1). F, *Kp-39* colonization by treatment group and intestinal segment on d1. Data were pooled from two independent experiments of within-litter controls with sequential dose (*n* = 9) and simultaneous dose (*n* = 9). In all instances, *n* refers to the number of pups of either sex. Mann-Whitney test (B and E) or two-way ANOVA (C, F, and G): *P≤0.05; **P≤0.005; ***P≤0.0005; ****P≤0.0001; ns, not statistically significant.

### EcN 1917 probiotic efficacy is independent of priority effect

Priority effect, which describes the impact that a species has on an environment depending on its order of arrival in that environment, is well documented across various micro- and macroscopic species and ecosystems (28–31). The reduction of *Kp-39* colonization following supplementation with EcN 1917 was not dependent on the timing of probiotic administration (Fig. 1D-G), indicating that there was no priority effect in this case. Littermates dosed sequentially with EcN 1917 followed by *Kp-39* challenge showed no significant difference in either total (Fig. 1E) or segmental (Fig. 1F) intestinal *Kp-39* colonization burden compared to those simultaneously dosed with EcN 1917 and *Kp-39* (Fig.1D). Further, there was no difference between the two dosing regimens in the ability of EcN 1917 to colonize any of the intestinal segments (Fig. 1G).

### Luminal oxygen content decreases over the first two weeks of normal development

Susceptibility to the development of neonatal dysbiosis and sepsis correlates with degree of prematurity and decreases with age following birth. In our model of *Kp-43816* sepsis, there is a natural resistance to the development of sepsis that emerges by p14 (15). In both human studies and in this murine model, increased developmental age correlates to an increase in the abundance of anaerobic bacterial species within the intestinal lumen (15).

Direct measurement of intestinal oxygen concentration in living animals, especially in animals as small as neonatal mice, is currently impractical (32). To better understand the luminal oxygen dynamics over this developmental window in the neonatal intestine, we therefore turned to a competitive index assay largely popularized by the Baumler and Winter groups (33–39). This assay compares the relative luminal growth of two isogenic bacterial strains which differ only in their ability to utilize a specific respiratory pathway, thereby allowing them to act as living biosensors for the environmental conditions of a given niche.

Enterobactericeae are uniquely suited for this role as they display a large degree of metabolic diversity. For the respiration of oxygen, *Escherichia coli* possess 3 cytochrome oxidases: CydA, AppC and CyoABCD, each with a different oxygen affinity (40). CydA is a high affinity oxidase (41), whereas CyoABCD is a lower affinity oxidase (41, 42). While relatively little is known about AppC, it is presumed to have an affinity which falls between the other two (40). While *E. coli* possess multiple reductases which allow for anaerobic respiration of a variety of substrates (*e.g.* nitrate, nitrite, fumarate, DMSO, *etc.*)(43–46) all of those pathways require molybdopterin cofactor, the synthesis of which requires the MoaA protein (47, 48). It is therefore straightforward to generate mutant strains of *E. coli* that are unable to respire at high O_2_ concentrations (Δ*cyoABCD* mutants), low O_2_ concentrations (Δ*cydA* mutants), or unable to carry out anaerobic respiration (Δ*moaA* mutants). In the experiments reported here, we used mutants of EcN 1917 which were generated for this purpose by Sebastian Winter (33). Utilizing this metabolic diversity to first examine the luminal conditions present at our timepoint of interest (p5, *i.e.* the fifth day after birth), we found that wild-type EcN 1917 demonstrated a significant growth advantage compared to the high-affinity Δ*cydA* deletion mutant, whereas no difference was seen in comparison to Δ*appC*, Δ*cyoABCD*, or Δ*moaA*, indicating the presence of a microaerophilic luminal environment in the p5 intestine (Fig. 2A).

**Figure 2.**
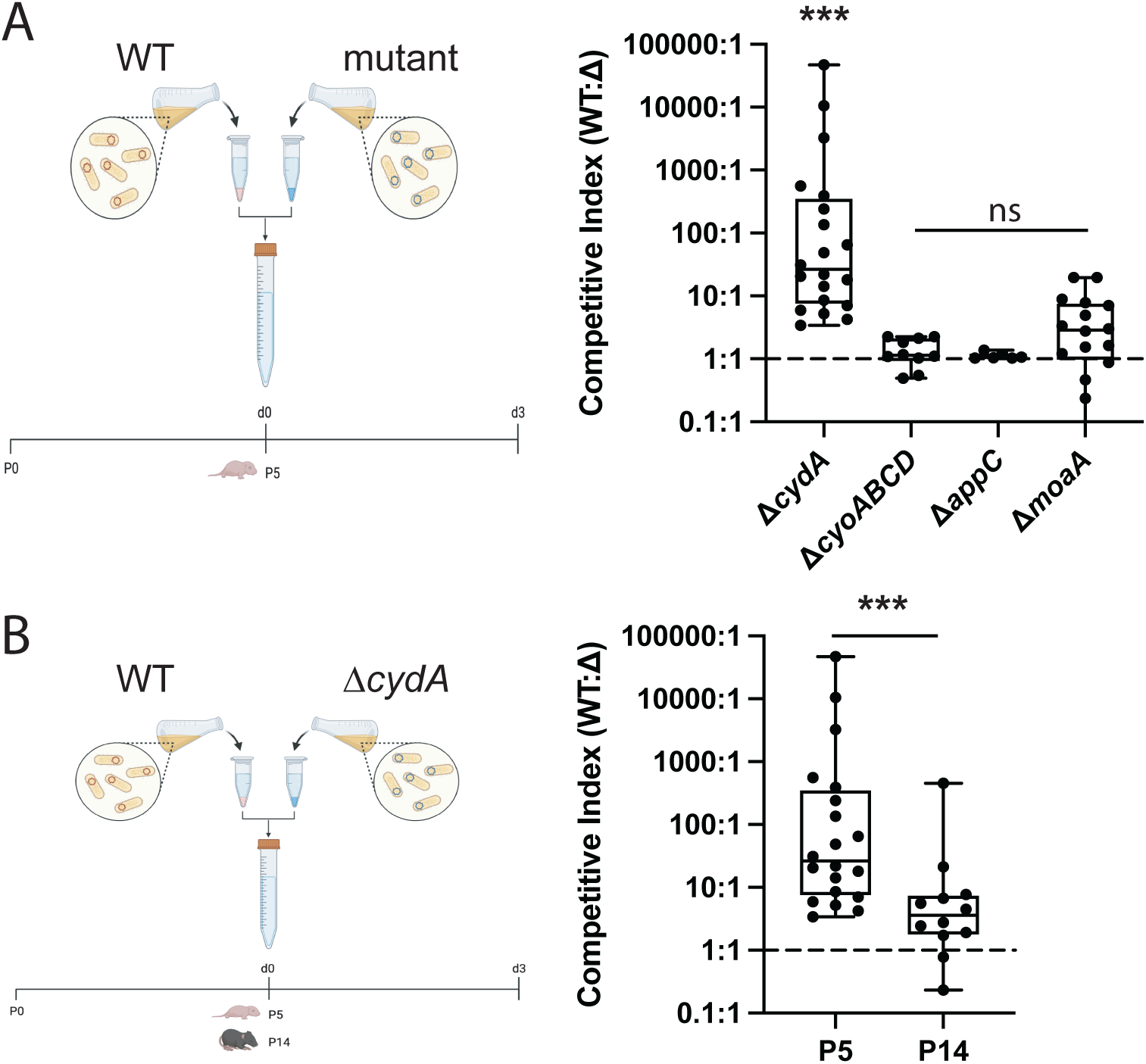
The p5 intestinal lumen is a microaerophilic environment and colonic luminal oxygen levels decrease over the first two weeks of normal development. A, p5 SPF mice (N = 6-20) were inoculated with a 1:1 mixture of *E. coli* wild type (WT and *cydA*, *cyoABCD*, *appC* or *moaA* mutants. The competitive index (CI; CFU of wild-type / CFU of mutant) was determined 3 days after inoculation. B, p5 and p14 SPF mice (*n* = 16-20) were inoculated with a 1:1 mixture of EcN 1917 wild-type (WT) and cydA mutant (ΔcydA). The competitive index (CI) was determined 3 days after inoculation. Mann-Whitney test: ***P≤0.01.

The competitive advantage conferred to wild-type EcN 1917 was abrogated by p14 (Fig. 2B), indicating a transition in the luminal oxygen status towards an anaerobic state, which is perhaps expected given the concurrent increase in colonization by obligate anaerobes previously described at this timepoint (15). Coupling this transition in luminal oxygen by p14 to the observation that there is a natural resistance to the development of dysbiosis and fulminant sepsis by this timepoint, it follows that a reduction in luminal oxygen may play a role in the protection against the development of dysbiosis.

### Deletion of the high-affinity oxidase CydA reduces EcN 1917 probiotic effect against *Kp39* dysbiosis in the distal small intestine and colon

Of course, in addition to its use as a biosensor of intestinal oxygen content, EcN 1917 is a potent probiotic in our model system (Fig. 1) but the extent to which EcN 1917’s respiratory activity contributed to inhibition of *Kp-39* colonization was unclear. In adult mice, respiration by EcN 1917 is required for inhibition of pathogenic facultative anaerobes, presumably by competition with those pathogens for respiratory substrates (33, 38, 39). In order to examine the role of oxygen respiration using the high-affinity CydA oxidase in the probiotic effect of EcN 1917 against *Kp-39* colonization in the neonatal intestine, littermate pups were administered either wild-type or Δ*cydA* EcN 1917 before challenge with *Kp-39* (Fig. 3A). Treatment with EcN 1917 Δ*cydA* resulted in higher total intestinal *Kp-39* colonization burden compared to treatment with wild-type EcN 1917 (Fig. 3B), indicating a role for oxygen respiration in probiotic action, however, this increase was not global but rather specifically due to increased *Kp-39* colonization in the distal small intestine and colon (Fig. 3C). There was no difference between the ability of wild-type and Δ*cydA* EcN 1917 to colonize any of the intestinal segments (Fig. 3D). Treatment with the Δ*cydA* mutant still resulted in total intestinal *Kp-39* colonization which was slightly lower than what was observed in the PBS vehicle treatment control alone (Fig. 3B). The ability of EcN 1917 to reduce *Kp-39* colonization cannot, therefore, be fully explained by competition between these two facultative anaerobes for oxygen.

**Figure 3.**
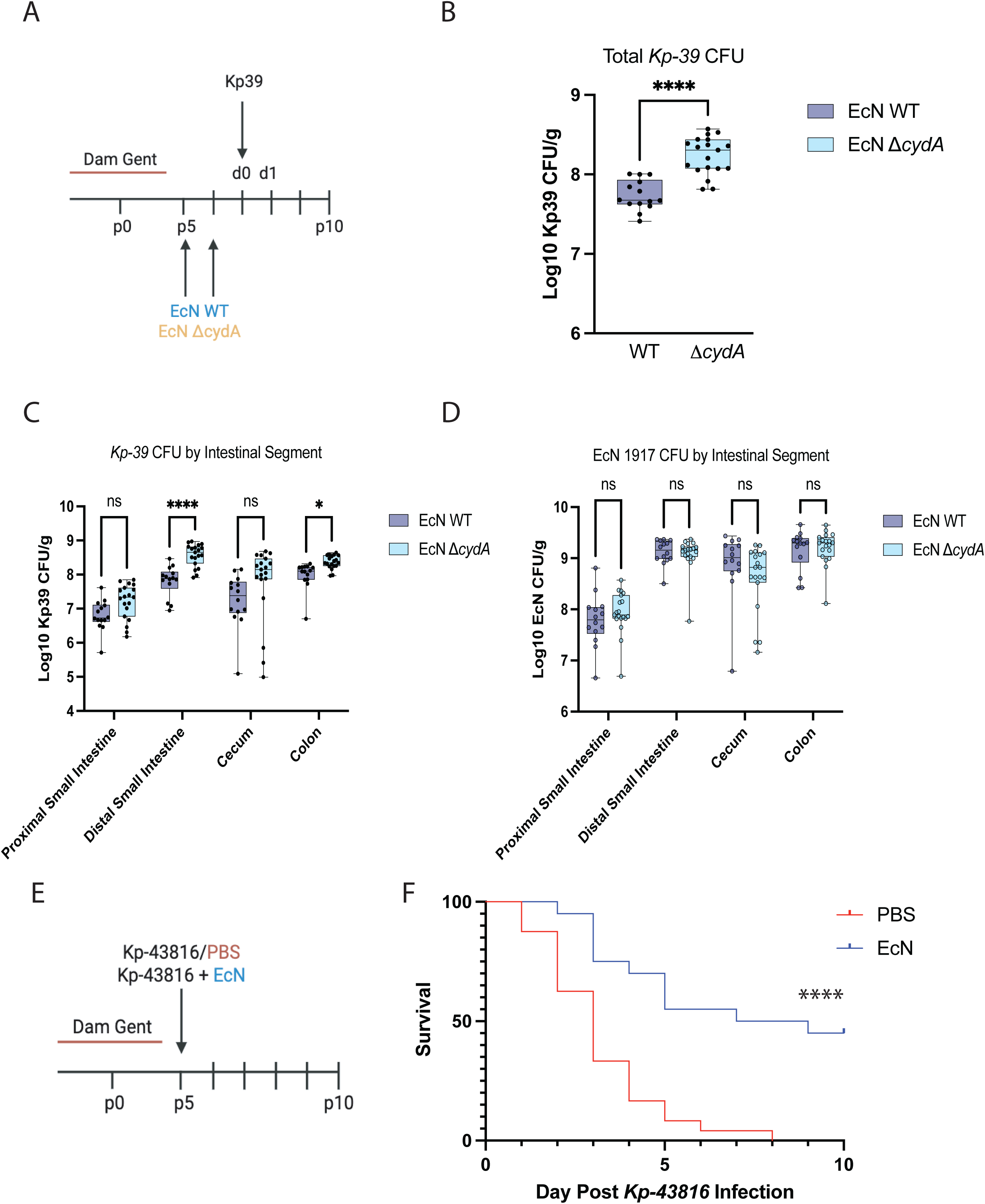
Probiotic effect of EcN 1917 is partially dependent on its ability to respire oxygen using its high affinity cytochrome oxidase CydA and extends survival against *Kp* sepsis when administered as a probiotic on P5. A, Schematic illustration of neonatal probiotic model. Littermate pups from gentamicin-treated dams were treated i.g. on day 5 (p5) and 6 (p6) of life with either EcN 1917 WT or EcN 1917 ΔcydA before infection with 10^7^ CFU *Kp-39* on day 7 (p7). B, Total intestinal *Kp-39* colonization burden by treatment group on treatment day 1 (d1). C, *Kp-39* colonization by treatment group and intestinal segment on d1. D, EcN 1917 colonization by treatment group and intestinal segment on d1. Data were pooled from three independent experiments of within-litter controls with WT (*n* = 14) and Δc*ydA* (*n* = 19). In all instances, *n* refers to the number of pups of either sex. Mann-Whitney test (B) or two-way ANOVA (C and D): *P≤0.05; **P≤0.005; ***P≤0.0005; ****P≤0.0001; ns, not statistically significant. E, Dams were separated 1-2 days before delivery and drinking water was supplemented with gentamicin (gent) until p4. On p5, pups were administered 10^7^ CFU *Kp-43816* with 10^7^ CFU EcN 1917 or without probiotic as a PBS negative control and monitored daily for sepsis. F, Kaplan-Meier curve. For survival analysis, data were pooled from multiple independent experiments with PBS (*n* = 24) and EcN (*n* = 20). In all instances, *n* refers to the number of pups of either sex. Log-rank (Mantel-Cox) test: ****P≤0.0001.

### EcN 1917 supplementation extends survival in *K. pneumoniae* sepsis

Given the established role of the microbiome in mitigating risk and mortality of neonatal sepsis through the initial prevention of dysbiosis (15), we sought to understand the extent to which our findings in the dysbiotic *Kp-39* model could be extrapolated to the prevention of fulminant sepsis using the more virulent *K. pneumoniae* strain *Kp-43816*.

Previous investigation of the impact of probiotics on prevention of sepsis by virulent *Kp-43816* were complicated by litter mortality following our standard probiotic model (15), which requires three separate gavages: two sequential probiotic doses followed by challenge with the virulent *Kp-43816*. This mortality was most likely due to direct bloodstream infection resulting from gavage-mediated microtraumas in the setting of such a virulent pathogen, and therefore probably does not reflect the situation *in vivo* where pathogens are thought to require transit into the bloodstream through the intestinal epithelium (49). However, given that equivalent reduction of *Kp-39* colonization was observed with simultaneous and sequential EcN 1917 / *Kp-39* dosing (Figs. 1E, F), the sepsis model was adapted to minimize trauma and consolidate treatment into a single gavage.

To this end, litters of gentamicin-reared pups were administered *Kp-43816* simultaneously with either EcN 1917 or a PBS negative control and monitored daily for mortality (Fig. 3E). Strikingly, probiotic supplementation with EcN 1917 significantly (P<0.0001) extended survival compared to PBS negative control (Fig. 6B), with 50% of pups surviving 10 days (as opposed to 100% mortality in pups without EcN 1917 treatment)(Fig. 3F).

### *L. murinus* V10 reduces *Kp-39* colonization burden across the intestinal span

*Ligilactobacillus murinus* is a well-known intestinal commensal in mice and other rodents (15, 50). We have previously shown that probiotic administration of the *L. murinus* V10 strain confers protection against the development of *Kp-39* dysbiosis in the neonatal murine intestine and is associated with prevention of late-onset sepsis in the same population (15). As expected, probiotic supplementation with *L. murinus* V10 (Fig. 4A) decreased total intestinal *Kp-39* colonization (Fig. 4B), which was due to a global decrease in colonization burden across each of the four examined intestinal segments (the proximal and distal small intestine, the cecum, and the colon) compared to a PBS control (Fig. 4C). Consistent with our observations of *L. murinus* V10, diverse *Lactobacillus* spp. are known to be able to colonize the entire length of the intestine (51, 52), however, unlike the obligate anaerobes that typically dominate the colon, lactobacilli are efficient colonizers of the small intestine (53), so it is perhaps unsurprising that the largest and most significant changes in *Kp-39* colonization following probiotic intervention with *L. murinus* V10 is seen in the proximal segments.

**Figure 4.**
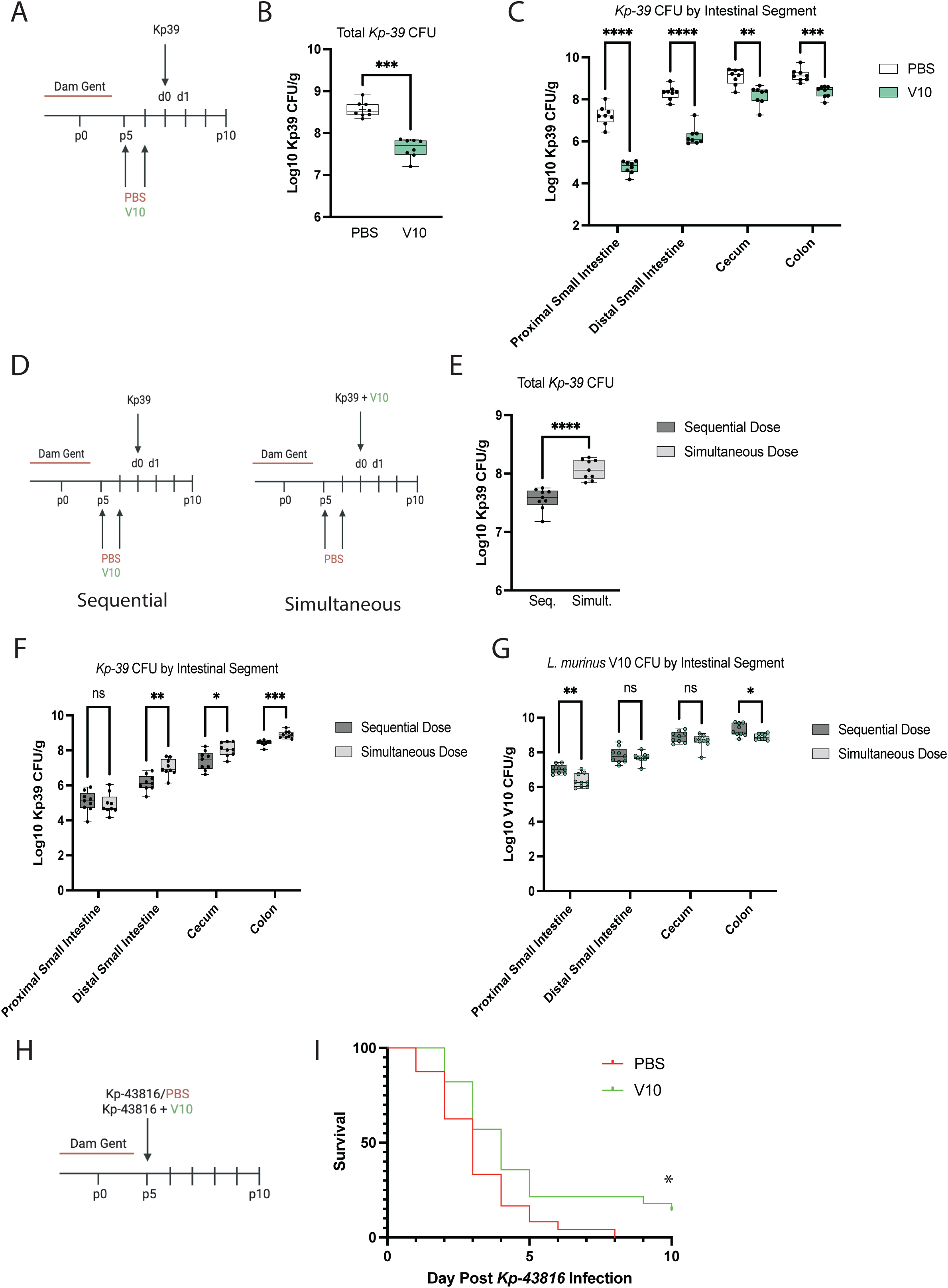
Biogeography and priority effect of *L. murinus* V10 probiotic effect in the neonatal intestine. A, Schematic illustration of neonatal probiotic model. Littermate pups from gentamicin-treated dams were treated i.g. on day 5 (p5) and 6 (p6) of life with either *L. murinus* V10 or PBS control before infection with 10^7^ CFU *Kp-39* on day 7 (P7). B, Total intestinal *Kp-39* colonization burden by treatment group on treatment day 1 (d1). C, *Kp-39* colonization by treatment group and intestinal segment on d1. Data were pooled from two independent experiments of within-litter controls with PBS (*n* = 8) and V10 (*n* = 8). D, Schematic illustration of neonatal priority effect model. Littermate pups from gentamicin-treated dams were treated i.g. on day 5 (p5) and 6 (p6) of life with either *L. murinus* V10 (sequential dose) or PBS control (simultaneous dose) before infection with 10^7^ CFU *Kp-39* (sequential dose) or 10^7^ CFU *Kp-39* and *L. murinus* V10 (simultaneous dose) on day 7 (P7). E, Total intestinal *Kp-39* colonization burden by treatment group on treatment day 1 (d1). F, *Kp-39* colonization by treatment group and intestinal segment on d1. H, *L. murinus* V10 colonization by treatment group and intestinal segment on d1. Data were pooled from two independent experiments of within-litter controls with sequential dose (*n* = 9) and simultaneous dose (*n* = 9). G, Dams were separated 1-2 days before delivery and drinking water was supplemented with gentamicin (gent) until p4. On p5, pups were administered 10^7^ CFU *Kp-43816* a with 10^7^ CFU *L. murinus* V10 or without probiotic as a PBS negative control and monitored daily for sepsis. H, Kaplan-Meier curve. For survival analysis, data were pooled from multiple independent experiments with PBS (*n* = 24) and V10 (*n* = 28). In all instances, *n* refers to the number of pups of either sex. Mann-Whitney test (B and E), two-way ANOVA (C, F, and G), or Log-rank (Mantel-Cox) test (H): *P≤0.05; **P≤0.005; ***P≤0.0005; ****P≤0. 0001; ns, not statistically significant.

### Prior exposure to *L. murinus* V10 enhances its probiotic effect

To assess the impact of priority effect on the ability of *L. murinus* V10 to reduce *Kp-39* colonization burden, littermate pups were administered the probiotic using either the normal dosing schedule (sequentially) or simultaneously with the *Kp-39* infectious challenge (Fig. 4D). In contrast to what we observed with EcN 1917 (Fig. 1), sequential dosing with *L. murinus* V10 resulted in lower total intestinal *Kp-39* burden compared to simultaneous dosing (Fig. 4E). This more precisely mapped to decreased *Kp-39* colonization in the distal small intestine, cecum and colon (Fig. 4F). *Kp-39* colonization in the proximal small intestine was not impacted by the timing of *L. murinus* V10 introduction.

Interestingly, while the timing of administration did impact colonization by *L. murinus* V10 (Fig. 4G), the intestinal segments with decreased colonization of *L. murinus* V10 (the proximal small intestine and colon) did not necessarily correlate to those with increased *Kp-39* microbial load (the distal small intestine, cecum, and colon).

Importantly, however, the simultaneous dosing schedule still reduced *Kp-39* colonization burden to a greater degree than what was seen with PBS treatment alone (refer to Fig. 4B), indicating that regardless of order of addition, *L. murinus* V10 is an effective probiotic that was able to reduce *Kp-39* colonization.

### Simultaneous *L. murinus* V10 supplementation increases survival during *K. pneumoniae* sepsis challenge

The priority effect observed for *L. murinus* V10 (Figs. 4D-G) complicated analysis of how well it protected against *Kp-43816* sepsis. However, given that a greater reduction of *Kp-39* colonization was observed with simultaneous probiotic / *Kp-39* dosing (Fig. 4E) than in PBS controls alone (Fig. 4B), we consolidated treatment with virulent *Kp-43816* into a single gavage with *L. murinus* V10 as well. Litters of gentamicin-reared pups were administered *Kp-43816* simultaneously with either *L. murinus* V10 or a PBS negative control and monitored daily for mortality (Fig. 4H). Probiotic supplementation with *L. murinus* V10 slightly, albeit significantly (P = 0.011) extended survival compared to the PBS negative control (Fig. 4I), albeit to a much lesser degree than EcN 1917 (Fig. 3F).

### Potential mechanisms of *L. murinus* V10 probiotic action

While *L. murinus* V10 was able to colonize all segments of the neonatal mouse intestine, it exerted its strongest inhibitory effects on *Kp-39* in the proximal segments (Fig. 4C), where oxygen is most abundant (54). Using EcN 1917 as a biosensor, we showed that the neonatal intestine contains substantial oxygen at early time points (Fig. 2), when these animals are susceptible to dysbiosis. In combination with our previous results showing a correlation between the ability of anaerobic bacteria to colonize and increased signal from a hypoxia-sensing probe in the neonatal mouse intestine with protection from dysbiosis (15), this led to us to hypothesize that oxygen-dependent responses might also help to explain the probiotic effect of *L. murinus* V10.

Supporting this hypothesis, we observed a striking phenotypic difference between *L. murinus* strain V10 and the non-protective *L. murinus* type strain ATCC 35020 (15). When grown in the presence of oxygen, *L. murinus* V10 aggregates and forms a dense, glutinous cell pellet (Fig. 5A) This does not occur in anaerobically-grown cells or in cultures of *L. murinus* ATCC 35020. Transcriptomics of *L. murinus* V10 in the presence and absence of oxygen revealed substantial changes in gene expression across the genome (815 out of 2,484 annotated protein-coding genes were significantly different in cells grown with or without oxygen)(Supplemental Data Set 1), including upregulation of several YSIRK-type signal peptide-containing cell surface proteins that could be involved in aggregation or adhesion (55–57) (Fig. 5B, Supplemental Tables S1-S3).

**Figure 5.**
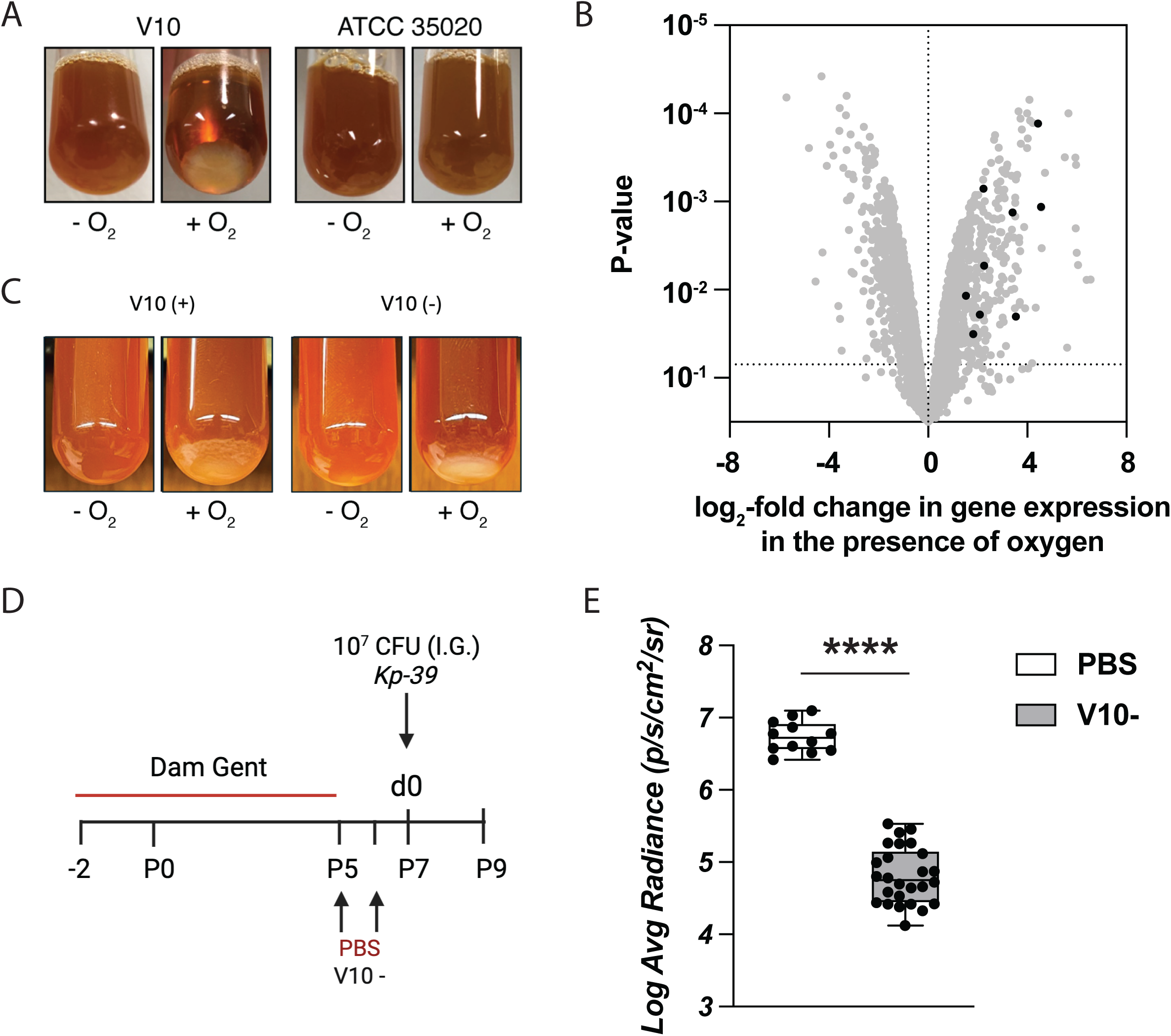
Megaplasmid pV10 is not responsible for the probiotic phenotype of *L. murinus* strain V10. A, *L. murinus* strains V10 and ATCC 35020 were incubated for 5h at 37°C in MRS medium either aerobically or anaerobically, then mixed by manual agitation. B, *L. murinus* V10 was grown to A_600_ = 2.0 in MRS medium either aerobically or anaerobically, then RNA was extracted and sequenced (n=3)(SeqCenter, LLC). For each gene, log_2_-fold change in average gene expression in aerobic cultures *versus* anaerobic cultures was plotted against the P-value for differential gene expression. Black dots indicate predicted highly expressed surface proteins (see Supplemental Table S3). C, *L. murinus* strains V10 (+) and *L. murinus* V10 (-) were incubated for 5 h at 37°C in MRS medium either aerobically or anaerobically, then mixed by manual agitation. D, Schematic illustration of neonatal probiotic model. Littermate pups from gentamicin-treated dams were treated i.g. on day 5 (p5) and 6 (p6) of life with either *L. murinus* V10 (-) or PBS control before infection with 10^7^ CFU *Kp-39* on day 7 (p7). E, *Kp-39* average radiance by treatment group (p/s/cm^2^/sr). Data were pooled from two independent experiments of within-litter controls with PBS (*n* = 10) and V10 (-) (*n* = 18). In all instances, *n* refers to the number of pups of either sex. Mann-Whitney test (E): ****P≤0.0001.

Neither transcriptional analysis nor comparative genomics can establish which gene or genes in *L. murinus* V10 are responsible for either aggregation or probiotic phenotypes. Unlike the readily manipulated EcN 1917, however, no molecular genetic tools have been reported for *L. murinus* to date, making experimental validation of such connections difficult.

Perhaps the most striking genetic element unique to *L. murinus* V10 in comparison to the *L. murinus* ATCC 35020 type strain, and certainly the easiest to address experimentally, is the presence of a 110 kbp megaplasmid in strain V10, eponymously termed “pV10” (GenBank Accession # CP040853.1). While many of the 110 genes on pV10 encode hypothetical proteins, there are also several homologs of sortase, adhesin, and S-layer proteins (58, 59), suggesting a possible role in cell-surface modification. We used novobiocin (60, 61) to cure *L. murinus* V10 of this megaplasmid, generating a plasmid-free derivative that is hereafter referred to as *L. murinus* V10 (-).

*L. murinus* V10 (-) demonstrated a similar degree of auto-aggregation in the presence of oxygen when compared to the *L. murinus* V10 parent strain harboring pV10 (V10 (+)), indicating that the pV10 plasmid is not required for this *in vitro* phenotype (Fig. 5C). Still, auto-aggregation and probiotic effectiveness are not necessarily actually linked, and the plasmid-cured *L. murinus* V10 (-) strain presented a unique opportunity to assess the impact of simultaneously knocking out 110 unique genes from *L. murinus* V10 on its ability to protect against *Kp-39* dysbiosis *in vivo* using the murine model of neonatal dysbiosis. To this end, littermate pups were administered either *L. murinus* V10 (-) or PBS negative control prior to challenge with *Kp-39*, using *Kp-39* luminescence as a rapid assessment tool to determine whether the plasmid-free strain had any ability to reduce *Klebsiella* burden (Fig. 5D). However, like *L. murinus* V10 (+)(15), *L. murinus* V10 (-) significantly reduced *Kp-39* luminescence compared to the PBS control (Fig. 5E), which indicated that pV10 was not required for the probiotic effect of *L. murinus* V10 against the dysbiotic expansion of *Kp-39* in the neonatal murine intestine.

### Dual Species Probiotic

Probiotics are often administered not as single bacterial strains, but in formulations containing multiple strains, species, and even genera (62–64). The rationale behind combining multiple probiotics of varying taxonomic relation is that protective effects may be complimentary; at a minimum additive, or in the best case synergistic. The evidence supporting this rationale is scanty. We therefore wanted to assess both whether and the extent to which combining our two probiotic species of interest impacted either the colonization *Kp-39* in the neonatal intestine or survival of *Kp-43816* sepsis relative to administration with single species probiotics alone.

### The effect of dual species (EcN 1917 and *L. murinus* V10) probiotic administration on *Kp-39* colonization is not additive and more closely mirrors *L. murinus* V10 supplementation

Littermate pups were administered either a dual-species probiotic treatment (10^7^ CFU of each) or single-species probiotic controls before challenge with *Kp-39* (Fig. 6). The dual-species probiotic significantly reduced total intestinal *Kp-39* colonization relative to both EcN 1917 (Fig. 6A) and *L. murinus* V10 (Fig. 6D) controls. Compared to EcN 1917 alone, administration of the dual-species probiotic was better able to reduce *Kp-39* colonization burden in both the proximal and distal small intestine and colon (Fig. 6B). In contrast, the only statistically significant reduction in *Kp-39* colonization burden by intestinal segment in dual-species relative to *L. murinus* V10 controls was in the distal small intestine (Fig. 6E). Importantly, there was no statistically significant difference in the intestinal colonization by either EcN 1917 or *L. murinus* V10 in single vs. dual species treated pups (Figs. 6C and F), indicating that neither probiotic strain affected colonization by the other.

**Figure 6.**
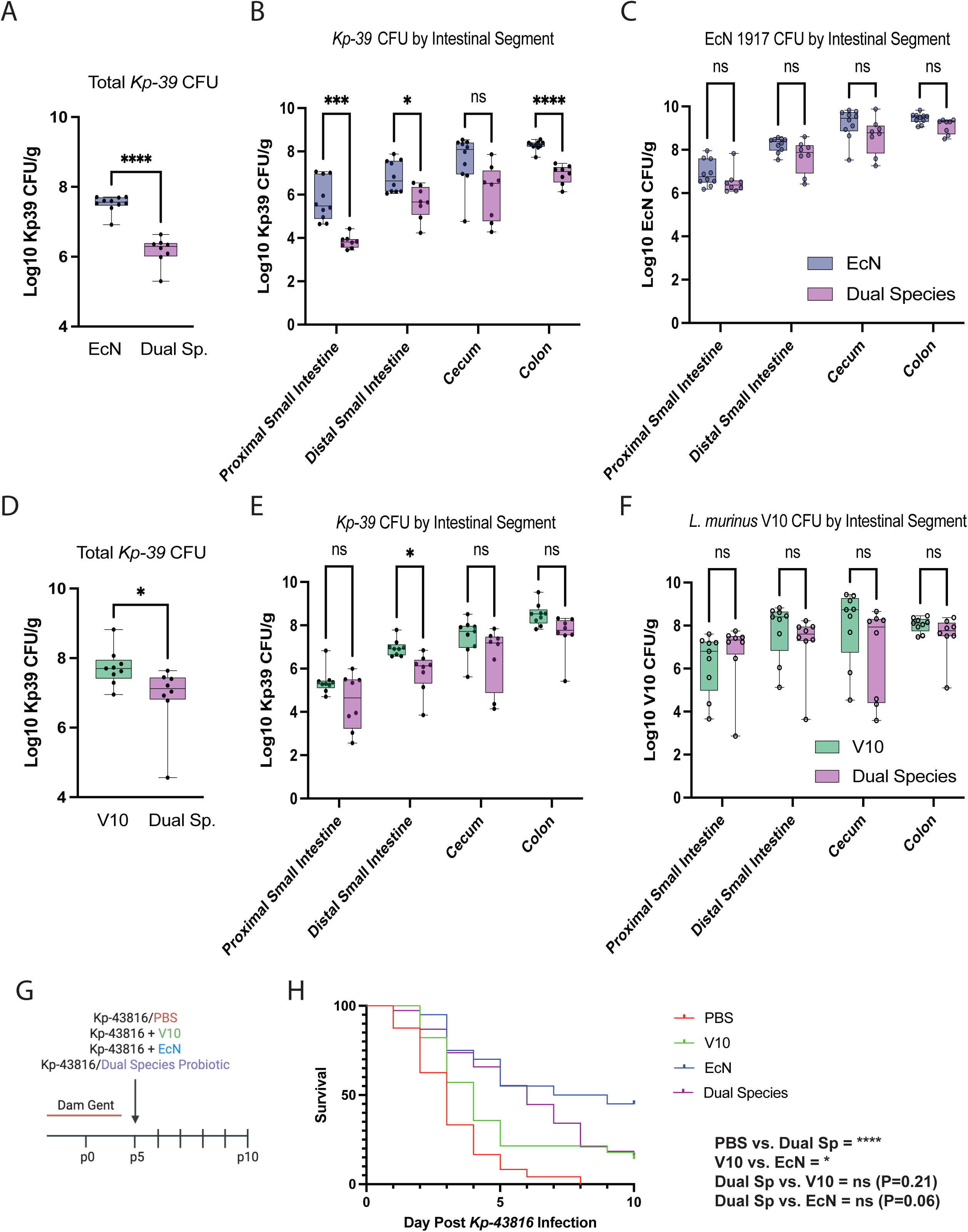
Biogeography of a *L. murinus* V10/EcN 1917 dual species probiotic effect in the neonatal intestine. A, Total intestinal *Kp-39* colonization burden by treatment group on treatment day 1 (d1). B, *Kp-39* colonization by treatment group and intestinal segment on d1. and intestinal segment on d1. C, EcN 1917 colonization by treatment group and intestinal segment on d1. D, Total intestinal *Kp-39* colonization burden by treatment group on treatment day 1 (d1). E, *Kp-39* colonization by treatment group and intestinal segment on d1. F, *L. murinus* V10 colonization by treatment group and intestinal segment on d1. Data were pooled from two independent experiments of within-litter controls with EcN (*n* = 10) and its dual species control (*n* = 8), V10 (*n* = 9) and its dual species control (*n* = 8). G, Dams were separated 1-2 days before delivery and drinking water was supplemented with gentamicin (gent) until p4. On p5, pups were administered 10^7^ CFU *Kp-43816* with 10^7^ CFU EcN 1917 or *L. murinus* V10 or without probiotic as a PBS negative control and monitored daily for sepsis. H, Kaplan-Meier curve. For survival analysis, data were pooled from multiple independent experiments with PBS (*n* = 24), EcN (*n* = 20), V10 (*n* = 28) and dual species (*n* = 38). In all instances, *n* refers to the number of pups of either sex. Mann-Whitney test (A and D) or two-way ANOVA (B, C, E, and F) or log-rank (Mantel-Cox) test (H): *P≤0.05; **P≤0.005; ***P≤0.0005; ****P≤0.0001; ns, not statistically significant.

### The effect of dual species (EcN 1917 and *L. murinus* V10) probiotic administration on survival of *K. pneumoniae* sepsis is not additive

To evaluate the impact of dual-species probiotic supplementation on survival against virulent *Kp-43816*, litters of gentamicin-reared pups were simultaneously administered either *Kp-43816* plus the dual species probiotic or *Kp-43816* with a PBS negative control and monitored daily for mortality (Fig. 6G). Dual-species probiotic supplementation significantly (P<0.0001) extended survival compared to the PBS negative control (Fig. 6H). Notably, however, survival analysis of litters administered the dual species probiotic formulation revealed no statistically significant difference from those administered either *L. murinus* V10 or EcN 1917 alone, indicating that there was not a strong additive effect on protection against death. Indeed, the final survival rate of mice administered the dual species probiotic in this experiment was identical to that of mice administered *L. murinus* V10 (20% survival) and much lower than that of mice administered EcN alone (50% survival).

## DISCUSSION

Understanding the factors underlying microbial community assembly in the neonatal intestine is critically important as we attempt to navigate the balance of promoting ecological conditions which favor accumulation of symbiotic microbes and prevent accumulation of dysbiotic microbes for both prevention and mitigation of disease development.

### The neonatal mouse as a model for probiotic ecology and mechanism

Currently, studies on probiotic efficacy and mechanism are typically carried out in adult animals or humans with complex, mature microbiota and have largely relied on indirect or relative readouts of intestinal bacterial burden (*e.g.* 16S or metagenomic DNA sequencing, analysis of fecal contents, or quantification of bacterial metabolites)(65, 66). In the work described here, the simplified community structure of the neonatal microbiome, coupled with baseline antibiotic “levelling,” and the delicate organismal size and tissue integrity of the neonatal mouse, which makes homogenization and quantification of the bacterial content of entire intestinal segments possible, afforded a unique opportunity to directly interrogate the ecological underpinnings of two probiotic species against the dysbiotic expansion of a clinically relevant pathogen.

### EcN 1917 prevents both *Kp-39* dysbiosis and fulminant *Kp-43816* sepsis

Overall, our results would suggest that EcN 1917 is a better overall probiotic in the neonatal intestine both for the prevention of dysbiosis and, especially, of fulminant sepsis. Of course, the use of a probiotic bacterium with such close genetic ties to pathogens in a highly vulnerable patient population such as premature infants raises important safety concerns. While there are very few reports of EcN 1917 causing extraintestinal infections in any human population (67, 68), there is no denying the substantial body of evidence linking pathogenic *E. coli* strains to a range of intra- and extra-intestinal infections and disorders (69–72). Notably particularly in the context of this work, *E. coli* is the one of the primary pathogenic species implicated in neonatal sepsis, albeit this is an association most strongly tied to early- and not late-onset sepsis (73–75).

One must also consider the long-term implications of persistent colonization at such an early developmental timepoint. Ecological succession describes the temporal changes in species composition of a given ecosystem over time (76). Pioneer species, defined as the initial populating members of nascent ecosystems (76), have the ability to significantly impact successional trajectories. This is particularly true in contexts with significant interspecies competition (77), such as the developing GI tract.

Considering successional theory in the context of the neonatal intestinal microbiome, it is therefore important to consider the potential for probiotics administered at this timepoint to act as pioneer species. While EcN 1917 was effective in the murine model we used at preventing immediate risk of dysbiosis and sepsis, persistent colonization may risk altering the succession events that otherwise define a healthy progression in the neonatal microbiome. Further studies are needed to assess the degree of persistence seen with EcN 1917 supplementation at this timepoint and what, if any, impact it has on normal microbial succession in the developing microbiome, and substantial safety evidence will be needed before EcN 1917 is likely to be approved for use in human infants, especially very vulnerable pre-term infants.

### Prevention of *Kp-43816* sepsis

While both *L. murinus* V10 and EcN 1917 were effective probiotics in the context of preventing the dysbiotic expansion of *Kp-39* throughout the intestinal span, they differed greatly in their ability to protect against the development of fulminant *Kp-43816* sepsis. However, as the *Kp-43816* model necessitated the simultaneous administration of the probiotic and *Kp-43816*, this discrepancy could be due in part to the relative sensitivity of *L. murinus* V10 to priority effect compared to EcN 1917. More work needs to be done to determine whether prior colonization with probiotics results in more effective sepsis prevention.

### More is not always merrier: variable effect of a dual species EcN 1917-*L. murinus* V10 probiotic

Combining *L. murinus* V10 and EcN 1917 into a single probiotic supplementation resulted in greater protection against the development of *Kp-39* dysbiosis but did not translate to increased protection against the development of fulminant *Kp-43816* sepsis. While the logic underlying multi-strain, multi-species, and even multi-genus probiotics seems sound at surface level, the fact of the matter is that combining multiple probiotic species does not necessarily equate to combining, much less synergizing, the impact of their individual probiotic effects. These are complex microbial communities, even in the simplified context of the neonatal intestine, and the biology of multi-species interactions are difficult both to study and predict.

### Mechanisms of probiotic action in the neonatal intestine

The mechanistic underpinnings of both probiotic species remain incompletely understood. The probiotic effect of EcN 1917 can at least in part be attributed to respiration using its high-affinity oxidase CydA, consistent with the current paradigm that oxygen and other respiratory substrates are important determinants of pathobiont growth in the intestine (78, 79). *L. murinus* V10 has unusual oxygen-dependent phenotypes *in vitro* and exerts strain-specific probiotic effects against *Kp-39* dysbiosis in the proximal intestine *in vivo*, both of which we were able to show were not driven by the 110-kb megaplasmid present in that strain. Further, dosing schedule of probiotics relative to *Kp-39* (priority effect) was found to play a role in *L. murinus* V10-mediated protection but did not impact the probiotic action of EcN 1917. We used transcriptomics to explore and identify potential mechanisms by which *L. murinus* V10 might exert its probiotic effects but have not yet established the molecular basis of those effects. One major factor limiting our ability to identify the molecular mechanism of V10’s probiotic activity is our very limited ability to genetically manipulate *L. murinus*. As of September 2025, a Google Scholar search did not identify any papers in which mutants of any strain of *L. murinus* were generated. We are currently working to develop tools for genetic manipulation of *L. murinus*, which will be necessary to further explore these questions.

### Conclusions

In total, we have demonstrated how two disparate probiotic species colonize the murine neonatal intestine and interact with a clinically relevant pathogen in the context of neonatal dysbiosis. Taken together, our data underscore the complex biological endeavor of probiotic intervention in the preterm intestine and the utility of applying community ecology as a framework for its analysis.

## MATERIALS AND METHODS

### Bacteria

#### Strains and plasmids used in this study

*E. coli* Nissle 1917 was isolated from a commercial probiotic product (Mutaflor)(80). *E. coli* Nissle 1917 *cydA*, *cyoABCD, appC,* and *moaA* mutants were a gift from Sebastian Winter (University of California - Davis)(81). *K. pneumoniae* strain *Kp-39* (aka Xen 39) was obtained from PerkinElmer. *K. pneumoniae* strain *Kp-43816* (aka ATCC 43816) was obtained from the American Type Culture Collection. *E. coli* and *K. pneumoniae* strains were transformed with low-copy plasmids pWSK29 (*bla*^+^) or pWSK129 (*kan*^+^)(82) to confer ampicillin or kanamycin resistance, respectively. *L. murinus* V10 was isolated from our mouse colony as previously described (15). *L. murinus* ATCC 35020 was obtained from the American Type Culture Collection. The identity of all strains was confirmed by whole-genome sequencing (SeqCenter, LLC), and the raw data are available on the NIH Sequence Read Archive (BioProject PRJNA979212).

#### Generation of rifampicin-resistant L. murinus

*L. murinus* V10 was incubated anaerobically at 37°C for 48 h in MRS media supplemented with 50 ug / mL rifampicin. Then, 100 uL of these cultures were spread on MRS plates supplemented with 25 ug/mL rifampicin and incubated anaerobically overnight at 37°C. Rifampicin-resistant colonies were isolated and genome sequenced by SeqCenter, LLC in Pittsburgh, PA to ensure no off-target mutations occurred, and the resulting strain MJG2601 (aka *L. murinus* V10-Rif^R^), containing an H490Y substitution in RpoB, was used in experiments mapping *L. murinus* CFU *in vivo*. No difference in growth was observed between wild-type and Rif^R^ derivatives of *L. murinus* (data not shown).

#### Plasmid curing

Plasmid curing was performed using novobiocin as a curing agent at varying concentrations from 0 to 30 μg/mL (60, 61). Briefly, *L. murinus* V10 was incubated anaerobically overnight in 5 mL MRS broth, then subcultured into MRS broth containing various concentrations of novobiocin and incubated anaerobically for 72 h at 37°C. After 72 h of incubation, 100 μL was plated onto MRS agar incubated overnight anaerobically at 37°C. Colonies were randomly selected and sent for genomic sequencing (SeqCenter, LLC) to confirm the absence of the pV10 plasmid, resulting in pV10-free strain MJG2665, which had no other mutations relative to wild-type *L. murinus* V10.

### Mice

C57BL/6J mice were obtained from JAX room RB09 then maintained at the University of Alabama at Birmingham (UAB). Colony husbandry and all experiments involving mice were approved by UAB’s Institutional Animal Care and Use Committee. For experiments with antibiotic-treated females, timed pregnant littermates were co-housed for the duration of their pregnancies without males until embryonic day (E) 19. Fresh hydropacs were weighed and gentamicin was added to a final concentration of 0.1 g/L. Pregnant females were gavaged i.g. with a loading dose of 5 mg gentamicin. Antibiotics remained in the drinking water until p5. 2 pregnant females who gave birth on the same day were co-housed with their pups through the remainder of the experiment. Pups were randomly assorted into groups before their first treatment.

### K. pneumoniae dysbiosis and sepsis infection model

*K. pneumoniae* were grown aerobically overnight in LB (10 g l^−1^ protease peptone 3, 5 g l^−1^ yeast extract, 10 g l^−1^ NaCl) at 37 °C and 250 r.p.m. The following day, the culture was diluted 1:50 in fresh LB and incubated for 2–3 h to reach exponential growth phase. Bacteria were washed twice with sterile PBS and resuspended to deliver 10^7^ CFU per 50-μl dose i.g. via a 22-gauge flexible polypropylene gavage needle (Instech). For each experiment, a test dose was serially diluted and plated to confirm between 5 × 10^6^ and 5 × 10^7^ CFU were delivered. Between each animal, the gavage needle was sanitized with 10% bleach and/or 70% ethanol, then dipped in sterile PBS for lubrication. Animals were imaged within 6 h of infection with the IVIS Lumina imaging system. Pups with luminescence signal in the thoracic cavity were euthanized and removed from the experiment. Litters were observed one to two times daily throughout the remainder of each experiment. Septic pups were removed immediately and clean bedding was provided daily to prevent cannibalism.

### Probiotic Supplementation

#### Sequential Dose Probiotic Model

Overnight anaerobic cultures of *Lactobacillus* strains grown in 5 mL of MRS broth were sub-cultured into 5 mL of fresh MRS broth then washed and diluted with sterile PBS. EcN 1917 was grown aerobically overnight in 5 mL of LB broth then sub-cultured in 5 mL of LB at 37°C at 200 r.p.m. A final dose of ∼10^6^ CFU per 50-μl dose for each organism was gavaged intragastrically on post-natal days p5 and p6. For each treatment, a test dose was serially diluted and plated on either MRS or LB agar, as appropriate, to confirm that between 1 x 10^5^ and 5 x 10^7^ CFU were delivered.

#### Simultaneous Dose Probiotic Model

Growth conditions for each organism were the same as noted above. A final dose of 1 x 10^7^ CFU per 50-μl dose for each organism was gavaged i.g. on P7. For each treatment, a test dose was serially diluted and plated on either MRS or LB agar to confirm between 5 × 10^6^ and 5 × 10^7^ CFU were delivered.

### Bacterial CFU determination

Complete intestinal segments were transferred to a pre-weighed microcentrifuge tube containing 1 ml of sterile PBS. Tissue was homogenized using a PowerGen500 homogenizer (Fisher), serially diluted in sterile PBS, and plated on LB agar containing 100 µg / mL ampicillin (EcN 1917) or LB agar containing 50 µg / mL kanamycin (*Kp-39*) and incubated overnight at 37 °C in ambient air or on LB agar containing 100 µg / mL ampicillin (EcN 1917 Δ*cydA*) or MRS agar containing 12.5 µg / mL rifampicin (V10-Rif^R^) and incubated overnight at 37 °C in a Coy anaerobic chamber in an atmosphere of 90% N_2_, 5% CO_2_, 5% H_2_. Dilutions resulting in between 30 and 300 bacterial colonies per plate were recorded for colony counts the following morning. During homogenization, the homogenizer was cleaned with 70% ethanol and sterile PBS between samples.

### Aggregation studies

*L. murinus* strains V10 and ATCC 35020 were incubated overnight at 37°C either aerobically or anaerobically in 5 mL MRS medium. The following morning, 100 µL of overnight culture was sub-cultured into 5 mL MRS medium either aerobically or anaerobically and incubated for 5 h at 37°C in, then mixed by manual agitation and visually assessed for aggregation.

### RNA Sequencing

For RNA extraction, single colonies of *L. murinus* V10 were inoculated into 5 mL of MRS and grown either anaerobically overnight at 37°C or aerobically at 37°C with shaking at 200 r.p.m. Cultures were then harvested, and RNA was extracted using the Qiagen RNeasy PowerMicrobiome Kit. mRNA sequencing was done by SeqCenter (Pittsburgh, PA). Samples were DNAse treated with Invitrogen DNAse (RNAse free). Library preparation was performed using Illumina’s Stranded Total RNA Prep Ligation with Ribo-Zero Plus kit and 10 bp unique dual indices (UDI). Sequencing was done on a NovaSeq X Plus, producing paired end 150 bp reads. Demultiplexing, quality control, and adapter trimming was performed with bcl-convert (v4.1.5). Quality control and adapter trimming was performed with bcl-con-vert. Read mapping was performed with HISAT2. Read quantification was performed using Subread’s featureCounts functionality. Read counts loaded into R were normalized using edgeR’s Trimmed Mean of M values (TMM) algorithm. Subsequent values were then converted to counts per million (CPM). Differential expression analysis was performed using edgeR’s glmQLFTest. Raw data are available on the NIH Sequence Read Archive (BioProject PRJNA979212).

## Data Availability

All raw data for this paper not deposited in the NIH Sequence Read Archive as described above are available on the FigShare data repository (https://doi.org/10.6084/m9.figshare.c.8038144).

Pre-publication reviewer access to these files is via the following links: https://dataview.ncbi.nlm.nih.gov/object/PRJNA979212?reviewer=5f8urvneu3g6acb3nof 72iuj41 https://figshare.com/s/74b3618e212fc6c3f10d.

## Acknowledgements

This work was supported by NIAID R01 AI164712 (to C.T.W. and M.J.G.) and NIGMS T32 GM008111 (to S.C.H.).

